# Differential mobility and self-association of Arc/Arg3.1 in the cytoplasm and nucleus of living cells

**DOI:** 10.1101/2021.12.02.470956

**Authors:** Per Niklas Hedde, Barbara Barylko, Derk D. Binns, David M. Jameson, Joseph P. Albanesi

## Abstract

Arc, also known as Arg3.1, is an activity-dependent immediate-early gene product that plays essential roles in memory consolidation. A pool of Arc is located in the postsynaptic cytoplasm, where it promotes AMPA receptor endocytosis and cytoskeletal remodeling. However, Arc is also found in the nucleus, a major portion being associated with promyelocytic leukemia nuclear bodies (PML-NBs). Nuclear Arc has been implicated in epigenetic control of gene transcription associated with learning and memory. In this study, we use a battery of fluorescence nanoimaging approaches to characterize the behavior of Arc in living cells. Our results indicate that in the cytoplasm, Arc exists predominantly as monomers and dimers associated with slowly diffusing particles. In contrast, nuclear Arc is almost exclusively monomeric and displays a higher diffusivity than cytoplasmic Arc. We further show that Arc moves freely and rapidly between PML-NBs and the nucleoplasm, and that its movement within PML-NBs is relatively unobstructed.

---

Arc (<u>A</u>ctivity-regulated <u>c</u>ytoskeleton-associated protein)^1^, also known as Arg 3.1 (<u>A</u>ctivity-regulated <u>g</u>ene 3.1)^2^, is a key regulator of synaptic plasticity and is required for the formation of long-term memories^3,4^. In response to neuronal activation, Arc protein levels increase rapidly (within minutes) in dendrites^5^, reflective of localized translation of pre-existing dendritic mRNAs, and more slowly (within hours) in the nucleus^6^. In spines, Arc promotes long-term depression (LTD) and synaptic weakening by facilitating endocytosis of AMPA-type glutamate receptors (AMPARs)^7,8^, but also enhances actin network formation^6^, a process more closely associated with long-term potentiation (LTP)^5^. In the nucleus, Arc plays a role in the regulation of gene transcription^6,9^, in part by facilitating histone acetylation^9^. We note that the ability of recombinant Arc to regulate transcription of genes involved in neuronal activity have been detected in heterologous cell systems, similar to those used in the present study^10^.

Arc self-associates into multiple oligomeric species *in vitro*, from dimers and tetramers up to 30-40-mers^11–14^. However, little is known about the oligomeric state of Arc in cells and essentially nothing is known about its behavior in the nucleus. To gain insight into these unresolved issues, we employed a combination of fluorescence-based nanoimaging approaches, including fluorescence fluctuation spectroscopy (FFS), Förster resonance energy transfer (FRET), and fluorescence recovery after photobleaching (FRAP). We conclude from our data that cytoplasmic Arc exists predominantly as a monomer-dimer equilibrium, whereas nuclear Arc is almost exclusively monomeric and diffuses more rapidly than its cytoplasmic counterpart. Moreover, although a major pool of Arc associates with promyelocytic leukemia nuclear bodies (PML-NBs)^15^, its movement between these membrane-less organelles and the surrounding nucleoplasm is rapid and apparently unobstructed within PML-NBs.

To measure the average diffusion coefficient of Arc in every region of the cell, we used the FFS approach of raster image correlation spectroscopy (RICS)^16^. As shown in Figure 1A-1C, the diffusion coefficients of Arc-EGFP ranged from ∼1-6 µm^2^/s in the cytoplasm of HeLa cells. By comparison, a diffusion coefficient of ∼28 µm^2^/s was reported for EGFP alone in the HeLa cell cytoplasm, also determined using RICS analysis^17^. The slow diffusion times of Arc-EGFP could indicate self-association into large particles or association with other cellular components, such as RNA, vesicles, and/or other proteins that significantly reduce its mobility. Interestingly, nuclear Arc displayed much faster diffusivity (∼3-12 µm^2^/s; Figure 1C-1E) than cytoplasmic Arc.

**Figure 1.**
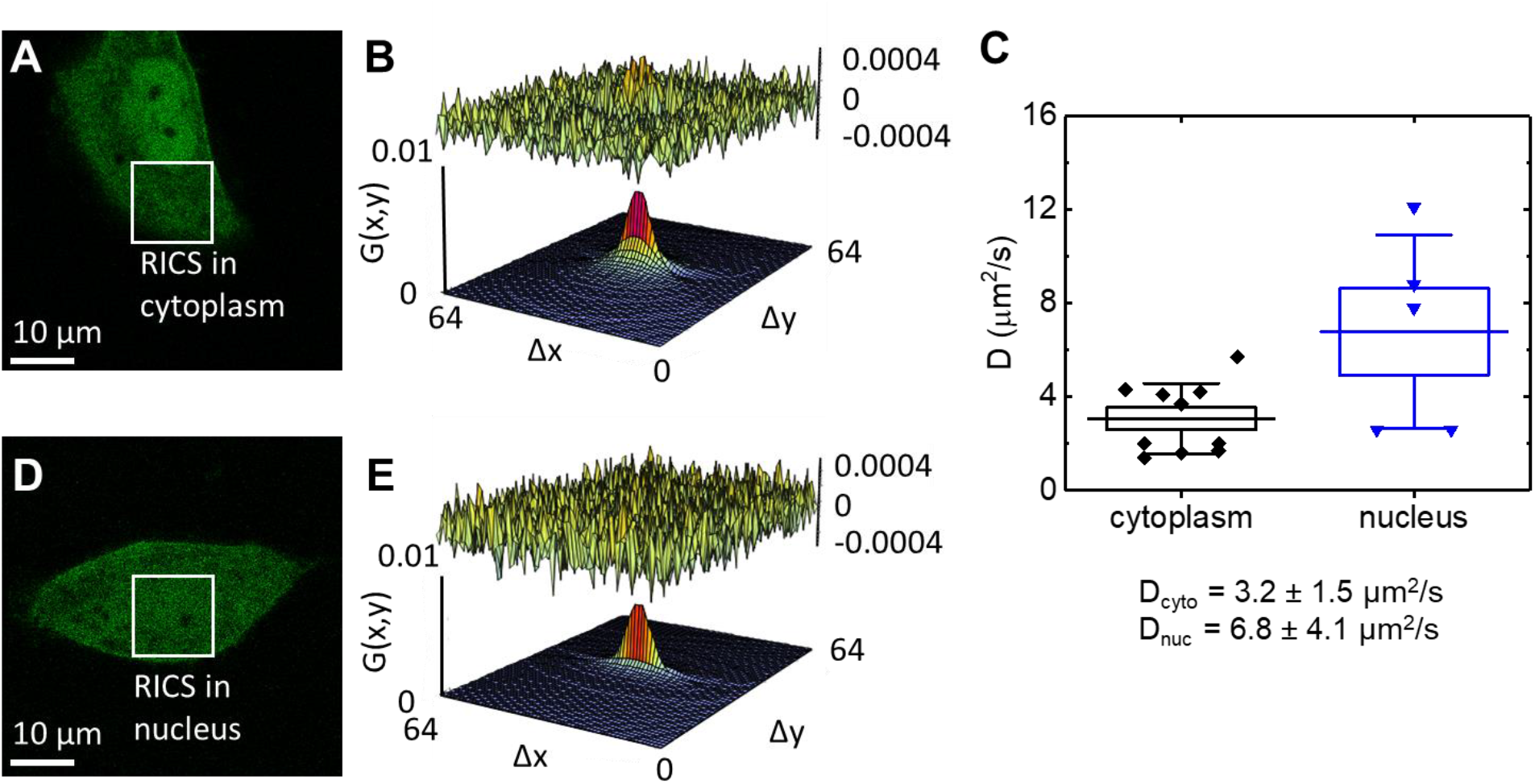
Measurement of Arc diffusion in cells. RICS analysis of Arc-EGFP in the cytoplasm (A,B) and nucleus (D,E) of HeLa cells. (A,D) Fluorescence images of representative cells expressing Arc-EGFP. (B,E) RICS analysis of the regions marked in (panels A and D). Fitted diffusion models are plotted and the residuals of the fits are displayed on top of the graphs. (C) Distribution of the measured diffusion coefficients (bar: mean, box: standard error (SE), whiskers: standard deviation (SD)). Number of measurements/cells analyzed: 9/5 (cytoplasm) and 5/5 (nucleus). Scale bars, 10 µm.

We next used the Number and Brightness (N&B) method to estimate the stoichiometry of EGFP-tagged Arc within moving particles in the cell. N&B utilizes the intensity fluctuations that occur when fluorescently labeled proteins pass through a small observation volume^18^. While the average intensity in two sample volumes containing the same number of fluorophores may be similar, the magnitude of the intensity fluctuations provides information about the protein stoichiometry. The measured molecular brightness can be normalized to a monomer standard, in this case the monomeric form of EGFP. In this manner, the stoichiometry of a labeled protein can be determined in every region of the cell. In addition, N&B measures the average number of particles within the observation volume, which can be converted to an average concentration when calibrated with the point spread function (PSF) volume. As shown in Supplementary Figure S1, the average expression level of EGFP-Arc was less than 300 nM in our experiments, a regime in which EGFP, which dimerizes with a KD of ∼100 μM^19^, does not aggregate. Figures 2A and 2B show, respectively, a representative fluorescence intensity image of an NIH-3T3 cell expressing EGFP-Arc and the brightness histogram of the same cell. Brightness ranges of monomers, dimers, trimers, and tetramers are indicated by green, orange, red, and purple boxes. Based on this color scheme, Figure 2C displays the brightness map of the entire cell. We quantified the stoichiometries separately in the cell cytoplasm and nucleus as shown in Figure 2D, where each point represents the average of a single cell. The brightness was significantly higher in the cytoplasm than in the nucleus. Recently, similar procedures were used to demonstrate that 1-2 copies of Arc incorporate into complexes on the plasma membrane of SH-SY5Y cells^20^.

**Figure 2.**
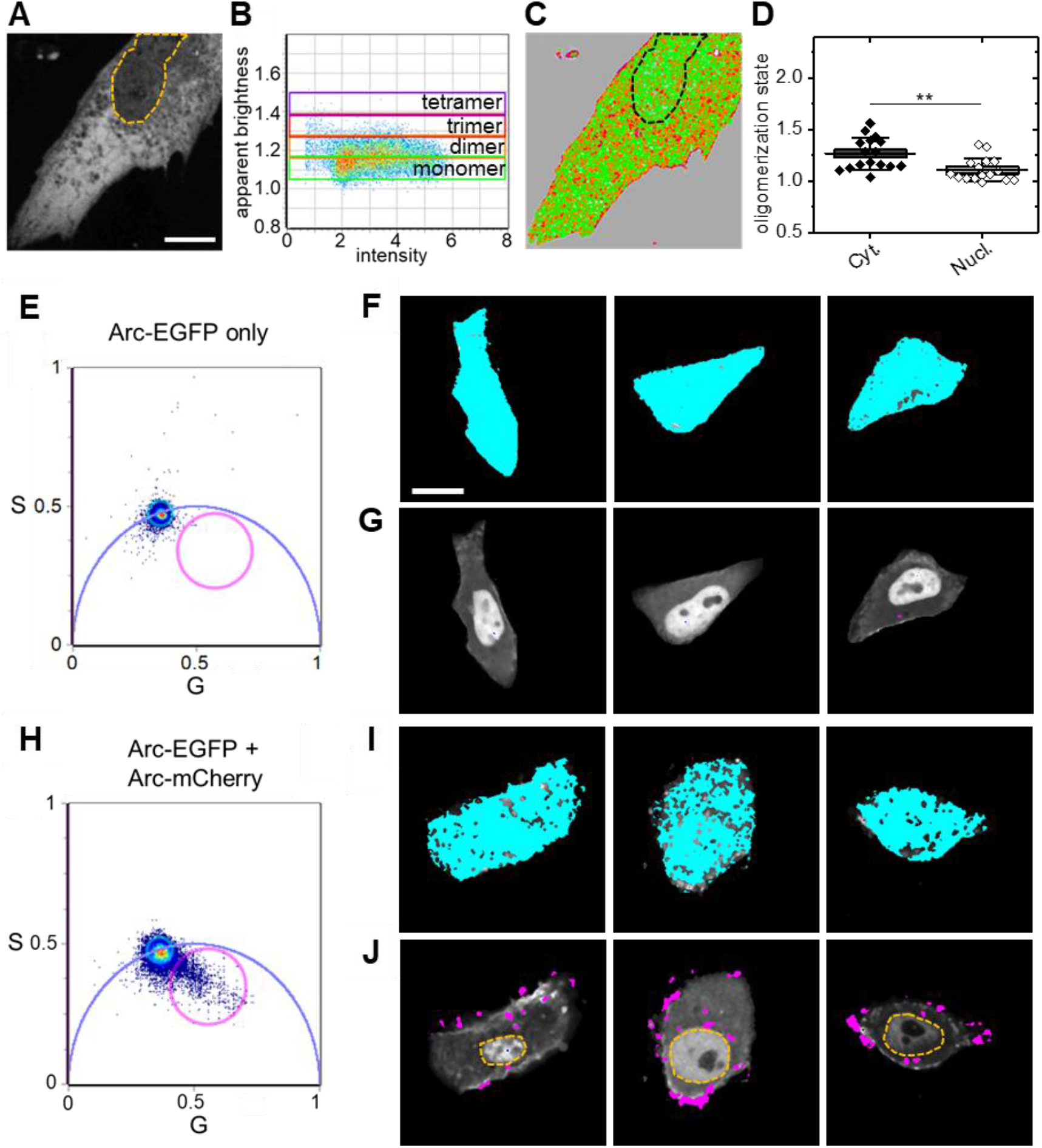
Oligomerization of cytoplasmic and nuclear Arc. (A-D) N&B analysis. Representative intensity image of an NIH-3T3 cell expressing EGFP-Arc. (B) Corresponding brightness plot of EGFP-Arc. Green, orange, red, and purple boxes indicate the brightness ranges of monomers, dimers, trimers, and tetramers. (C) Brightness map of the cell shown in panel A color-coded according to panel B. (D) Statistical analysis of the measured oligomerization states (stoichiometries) using the Mann-Whitney non-parametric test (bar: mean, boxes: ± SE, whiskers: ± SD, ** p < 0.01). Number of cells analyzed N = 14. (E-J) FRET/FLIM analysis. (E) Phasor plot pixel lifetime distribution of cells expressing Arc-EGFP (donor only). (F) Pixels corresponding to non-quenched EGFP were selected by the cyan cursor and highlighted in the fluorescence intensity images. (G) Pixels corresponding to quenched EGFP were selected by the magenta cursor and highlighted in the same fluorescence intensity images. (H) Phasor plot pixel lifetime distribution of cells co-expressing Arc-EGFP and Arc-mCherry. (I) Pixels corresponding to non-quenched EGFP were selected by the cyan cursor and (J) pixels corresponding to quenched EGFP were selected by the magenta cursor and highlighted in the fluorescence intensity images. Scale bars, 10 µm.

Taken together, the data presented in Figures 1 and 2A-2D demonstrate that 1-2 copies of Arc move together in the cytoplasm, whereas nuclear Arc is almost exclusively monomeric and displays a significantly higher mobility than cytoplasmic Arc. The striking difference between the diffusion coefficients of cytoplasmic and nuclear Arc cannot be explained by the less than two-fold difference in their stoichiometries, but instead suggests that Arc associates with larger, more slowly moving complexes in the cytoplasm than in the nucleus.

Although our N&B measurements revealed that, on average, 1-2 molecules of EGFP-tagged Arc move together in a complex, they do not unequivocally establish that the molecules exist as monomers and/or dimers within that complex. To resolve this issue, we turned to measurements of FRET, which can occur if an acceptor fluorophore comes within a few nanometers of a donor. Energy transfer causes a reduction in both the intensity and lifetime of the donor; however, lifetime measurements are generally a more robust reporter of FRET in living cells^21^. We expressed Arc-EGFP (the donor), either alone or together with Arc-mCherry (the acceptor) in HeLa cells and acquired lifetime images by pulsed 880-nm two-photon excitation. The fluorescence lifetime of each pixel was calculated and displayed using phasor plots^22^. Figure 2E shows the 2D histogram of the pixels of three images of HeLa cells that only expressed Arc-EGFP (donor only). The center of mass of the distribution falls on the universal semicircle, indicating a single exponential lifetime decay as expected for unquenched Arc-EGFP. In Figure 2F, these pixels are highlighted with a cyan color, showing their uniform distribution within the entire cell. As expected for this donor-only control, no quenched Arc-EGFP was detected, as demonstrated by the absence of pixels highlighted with a magenta color in the cells (Figure 2G). As an additional control, no energy transfer was detected when EGFP was expressed together with mCherry (Supplementary Figure S2). In contrast, phasor plots of data from cells co-transfected with Arc-EGFP and Arc-mCherry displayed a comet tail extending from the universal semicircle towards the zero-lifetime point (S = 0, G = 1) (Figure 2H). In addition to unquenched EGFP (cyan pixels in Figure 2I), magenta pixels representing quenched EGFP were evident in the corresponding images (Figure 2J). Interestingly, regions of high energy transfer were exclusively detected in the cell cytoplasm and at the plasma membrane; none were detected in the nucleus (outlined by the dashed yellow line in Figure 2J). Combining these results with the N&B data, we conclude that cytoplasmic Arc exists predominantly in a monomer-dimer equilibrium, whereas nuclear Arc tends to remain monomeric.

Prior studies indicated that a pool of Arc accumulates in promyelocytic leukemia nuclear bodies (PML-NBs)^9,15^, which are phase-separated membrane-less organelles involved in (among other functions) epigenetic control of gene transcription. The absence of FRET in the nucleus indicates that Arc does not oligomerize within PML-NBs (Figure 2J). However, it is possible that Arc in PML-NBs is relatively immobile and therefore undetectable by RICS and N&B, which rely on signal fluctuations caused by protein translocation between image pixels. To explore this possibility, we co-expressed Arc-mCherry and PML-EGFP in HEK293 cells and applied dual channel FRAP to individual PML-NBs. Figure 3A and 3B show pre-bleaching fluorescence images for PML-EGFP and Arc-mCherry, respectively. One to three PML-NBs were bleached per cell (arrows). Typical fluorescence recovery curves for each protein are plotted in Figure 3C and 3D. In general agreement with previous studies^23^, the average recovery time of PML-EGFP fluorescence was >100 s (Figure 3E), indicative of the relatively slow translocation of PML into and out of PML-NBs. In contrast, the average recovery time of Arc-mCherry fluorescence was only ∼10 s (Figure 3E), indicating a rapid exchange of Arc between the PML-NB-associated and non-PML-NB-associated pools of nuclear Arc. Further, we observed a lower level of bleaching for Arc-mCherry compared to PML-EGFP together with an overall reduction in the nuclear Arc-mCherry signal immediately after the 0.3-3 s bleaching period, indicating that diffusion of bleached Arc-mCherry away from the PML-NBs was so rapid that it could partially replace unbleached Arc-mCherry in the rest of the nucleus before monitoring of the recovery process began.

**Figure 3.**
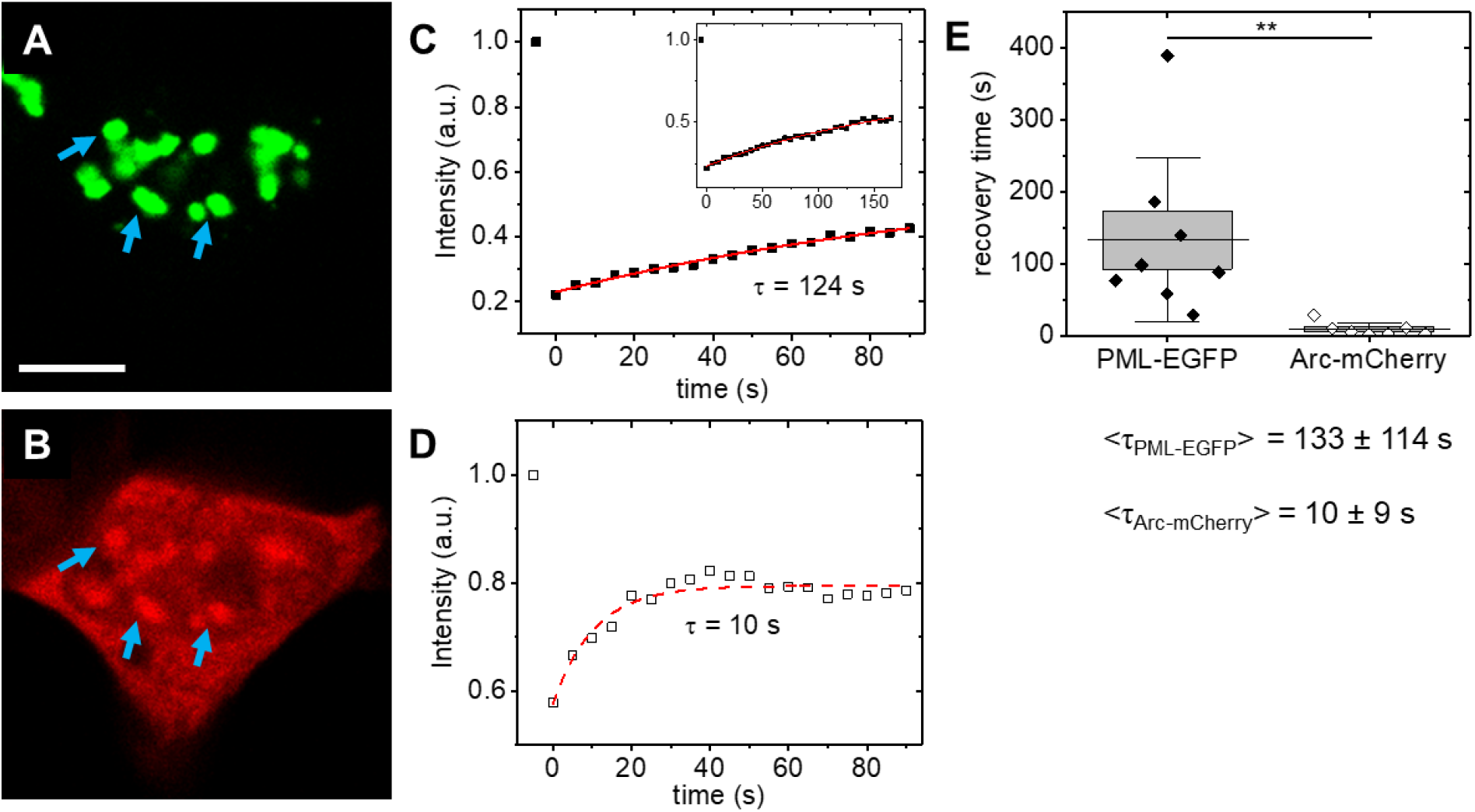
FRAP analysis of Arc in PML-NBs. (A,B) Pre-bleaching fluorescence images of PML-EGFP (A) and Arc-mCherry (B) in HEK293T cells. One to three PML-NBs were bleached per cell (arrows). (C,D) Representative fluorescence recovery data of PML-EGFP (C) and Arc-mCherry (D) fitted with a single exponential association model. The inset in panel C shows the recovery of PML-EGFP at longer times. (E) Statistical analysis of the measured fluorescence recovery times using the Mann-Whitney non-parametric test (bar: mean, boxes: ± SE, whiskers: ± SD, ** p < 0.01). The fluorescence recovery of N = 8 (PML-EGFP) and N = 7 (Arc-mCherry) PML-NBs was analyzed. Scale bar, 10 µm.

While FRAP demonstrated a rapid exchange of Arc protein between the nucleoplasm and PML-NBs, this single point method does not reveal the paths taken by the molecules. To visualize and quantify barriers to the motion of Arc in the nucleus, we applied the two-dimensional spatial pair correlation function (2D-pCF) approach^24^ (Figure 4). This approach measures the average time a molecule takes to move between two points in a cell by cross-correlating the intensity fluctuations at specific points in an image. Rapid acquisition of an image time series was followed by pair correlation of each pixel with neighboring pixels at equally spaced angles around the origin at the pair correlation distance, *δr* = 4 pixels. The resulting set of pCFs was then examined for asymmetries. If no obstacles are present within the pair correlation distance, the correlation function will be symmetric/equal in all directions. Near a barrier, where the motion of the molecule of interest is restricted or absent, the set of pCFs will be deformed. This directional preference (motion anisotropy) can be quantified by moment analysis (Figure 4G). For barrier mapping, Arc-mCherry and PML-EGFP were expressed in HEK293 cells. Representative fluorescence images of the nucleus are shown for PML-EGFP (Figure 4A and 4B) and for Arc-mCherry (Figure 4C and 4D). Rapid time sequences of Arc-mCherry were acquired and subjected to 2D-pCF analysis. To visualize how obstacles shape the effective flow of Arc molecules, a map was created by drawing lines with their length proportional to the motion anisotropy in each pixel on top of the PML-EGFP intensity image, where the line orientation and color represents the movement direction (Figure 4E and 4F). This map highlights areas of highly directional and confined Arc movement. As expected, high motion anisotropy was observed where PML-NBs interfaced with the nucleoplasm, as barriers are needed to maintain the concentration difference between compartments. To our surprise, we found that Arc movement was largely unobstructed within PML-NBs (Figure 4H). By comparison, barriers modified the motion of Arc in the nucleoplasm, with possible obstacles including heterochromatin structures. Together with the FRAP data indicating that the fluorescence recovery of Arc in PML-NBs was much faster than that of PML itself, we conclude that Arc is loosely associated with other PML-NB components.

**Figure 4.**
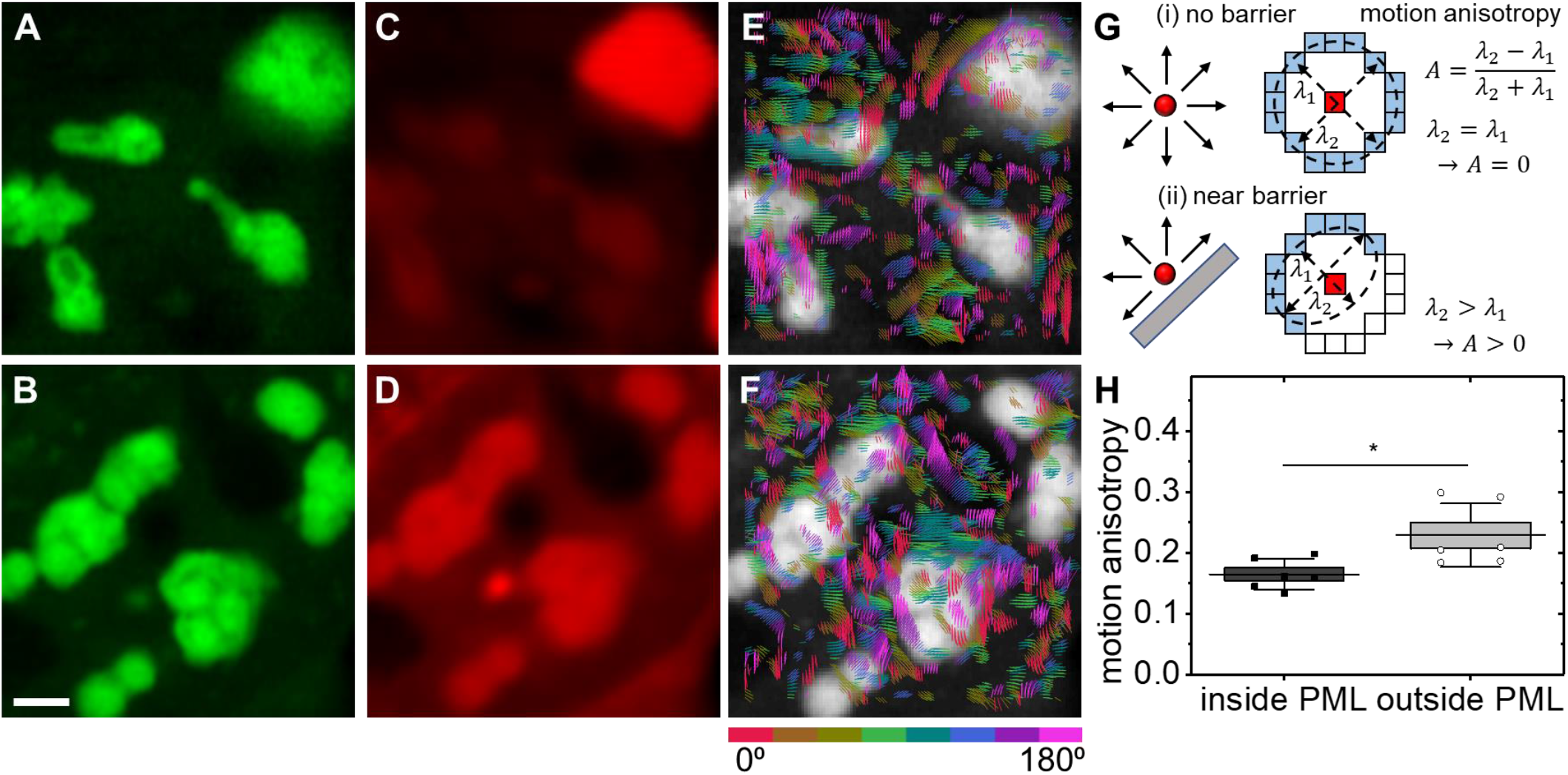
2D-pCF analysis of Arc in the nucleus. (A,D) Representative fluorescence images of PML-EGFP (A,B) and Arc-mCherry (C,D) in the nucleus of HEK293T cells. (E,F) Corresponding Arc-mCherry motion anisotropy maps (colored lines) overlaid with the PML-EGFP intensity images (grayscale). The length of each line corresponds to the motion anisotropy in each pixel; both the line orientation and color indicate the dominant direction of movement. For visualization purposes, only lines with anisotropies >0.15 (panel E) and > 0.25 (panel F) were plotted. (G) Illustration of the 2D-pCF principle. Without barriers at the pair correlation distance (here: 4 pixels, 444 nm at the sample) the set of pCFs is symmetric. Near a barrier, the molecule motion is delayed or inhibited, resulting in an asymmetric set of pCFs. The degree of motion distortion can be quantified with the motion anisotropy formula, where *λ*_*1*_ and *λ*_*2*_ are the short and long axes of each pCF set. (H) Statistical analysis of the measured motion anisotropy within PML-NBs and in the nucleoplasm (outside PML-NBs) using the Mann-Whitney non-parametric test (bar: mean, boxes: ± SE, whiskers: ± SD, * p < 0.05). Number of cells analyzed N = 6. Scale bar, 2 µm.

Evidence from the VanDongen group supports a model in which Arc recruits the Tip60 histone acetyltransferase complex to PML-NBs, triggering a chromatin remodeling program that regulates long-term memory formation^9^. However, the mechanisms that target Arc itself to PML-NBs remain to be identified. Many PML-NB components, including PML itself, are SUMOylated and/or contain SUMO-interaction motifs (SIMs). Although Arc is both SUMOylated and contains a putative SIM^25^, we were unable to detect a direct interaction with PML using fluorescence lifetime-based FRET (Supplementary Figure S3). Interestingly, Arc may not only function as a transcriptional regulator within PML-NBs, but may also contribute to their formation or stabilization, as elevation of Arc expression increases the number of PML-NBs in neurons^6^ and HEK293 cells^15^.

## METHODS

### Materials

Mouse Arc cDNA, cloning and mutagenesis reagents were from Thermo Scientific. Primers were from Invitrogen. Lipofectamine 3000 was from ThermoFisher.

### Expression of recombinant Arc

To generate fluorescent protein-tagged Arc for mammalian expression, GST-Arc was used as a template for subcloning into appropriate vectors. pEGFP and mCherry vectors were from Clontech (with CMV promoter) with either C1 or N1 cloning sites to obtain the tag either at the N-terminus or C-terminus of Arc, respectively. Cloning sites were XhoI and BamHI with stop codon for N-terminal tag or without stop codon for C-terminal tag. All DNA constructs were verified by sequencing.

### Cell culture

HEK293, HeLa, and NIH-3T3 cells were plated on fibronectin-coated dishes, cultured in DMEM (supplemented with 10% FBS and 1% Pen-Strep) and transfected with Lipofectamin 3000. They were imaged 15-25 hours after transfection either at room temperature (RICS, FLIM, N&B), or at 37° C and 5% CO2 (FRAP and 2D-pCF).

### Raster Image Correlation Spectroscopy (RICS)

HeLa cells transfected with Arc-EGFP plasmid were imaged with an Olympus FV1000 confocal microscope in photon counting mode. Fluorescence was excited at 488 nm and detected at 505-525 nm with a 60x, NA 1.2 water immersion lens. Per data set, 100 frames of 256 × 256 pixels were acquired with a 20 µs pixel dwell and 6.24 ms line scan time, sample pixel size was 50 nm, the waist, *w*0, of the point spread function was 270 nm. A moving average of 10 frames was subtracted to remove the immobile fraction. Data were analyzed in SimFCS (Globals Software, CA).

### Fluorescence lifetime imaging (FLIM)

NIH-3T3 cells transfected with Arc-EGFP and Arc-mCherry plasmids were imaged with a Zeiss LSM880 laser scanning microscope set up for FLIM. EGFP fluorescence was excited at 880 nm (two-photon excitation, 80 MHz) and detected at 510-560 nm with a 40x, NA 1.2 water immersion lens in non-descanned mode using a hybrid photomultiplier detector (HPM-100, Becker & Hickl, Germany) coupled to a FLIMBox (ISS, Champaign, IL). Per data set, 35 frames of 256 × 256 pixels were acquired with a pixel dwell time of 16 µs. Data were analyzed in SimFCS. Before FLIM, the presence of both donor (EGFP) and acceptor (mCherry) were verified with 488-nm and 594-nm excitation.

### Number and Brightness (N&B)

NIH-3T3 cells were transfected and imaged as described for RICS. Per data set, 100 frames of 256 × 256 pixels were acquired with a pixel dwell time of 10 µs; sample pixel size was 165 nm. For calibration, a solution of monomeric EGFP (500 nM) was prepared in PBS buffer. The true molecular brightness of EGFP monomers was measured as *b* = 0.11. All instrument parameters were kept identical. Data were analyzed in SimFCS (Globals Software, CA).

### Fluorescence Recovery after Photobleaching (FRAP)

HEK293T cells transfected with Arc-mCherry and PML-EGFP plasmids were imaged with a Zeiss LSM880 laser scanning microscope set up for FRAP. EGFP fluorescence was excited at 488 nm and detected at 510-560 nm, mCherry fluorescence was excited at 594 nm and detected at 610-680 nm, both with a 40x, NA 1.2 water immersion lens. Per data set, one pre-bleaching and 35 post-bleaching frames of 256 × 256 pixels were acquired at 5 s intervals (4 µs pixel dwell time, 4x line average). PML bodies were bleached in 1-3 circular regions per cell of 1-3 µm diameter for 0.1-1 s each. Recovery data were analyzed in Origin (OriginLab, MA), a single exponential association model was used, 19 data points were fitted for the rapid recovery of Arc-mCherry and 34 data points were fitted for the slower recovery of PML-EGFP.

### Two-dimensional pair correlation function (2D-pCF)

HEK293T cells were transfected and imaged as described for FRAP. The LSM880 was set up for fast acquisition with the Airy detector. Per 2D-pCF data set, 10,000 frames of 116 × 116 pixels were acquired at 50 frames per second. Data were analyzed in SimFCS (Globals Software, CA) as previously described^24^. The pair correlation distance was 4 pixels, the pixel size at the sample was 111 nm. Based on the PML-EGFP images, masks were drawn to separately quantify the motion anisotropy in PML-NBs and the nucleoplasm.

## Abbreviations

AMPA: α-amino-3-hydroxy-5-methyl-4-isoxazolepropionic acid
DMEM: Dulbecco’s Modified Eagle Medium
FBS: Fetal Bovine Serum

**Accession IDs** of proteins used in this study: mouse Arc/Arg3.1: UniProt ID:Q9WV31;

## AUTHOR INFORMATION

### Author Contributions

All authors have contributed to writing the manuscript. JPA, BB, and DMJ were responsible for the experimental design. BB and DDB performed cloning. PNH performed nanoimaging and analysis.

### Notes

The authors declare no competing financial interest.

## Acknowledgement

We thank Dr. Michael Rosen (UTSW) for PML-EGFP plasmids and Steve Stippec (UTSW) and Joel Villarreal (UTSW) for invaluable technical assistance. The nanoimaging experiments were performed at the Laboratory for Fluorescence Dynamics (LFD) at the University of California, Irvine (UCI). The LFD is supported jointly by the National Institute of General Medical Sciences of the National Institutes of Health (P41GM103540), and UCI.

## REFERENCES

(1) Lyford, G. L.; Yamagata, K.; Kaufmann, W. E.; Barnes, C. A.; Sanders, L. K.; Copeland, N. G.; Gilbert, D. J.; Jenkins, N. A.; Lanahan, A. A.; Worley, P. F. Arc, a Growth Factor and Activity-Regulated Gene, Encodes a Novel Cytoskeleton-Associated Protein That Is Enriched in Neuronal Dendrites. Neuron 1995, 14 (2), 433–445. https://doi.org/10.1016/0896-6273(95)90299-6.

(2) Link, W.; Konietzko, U.; Kauselmann, G.; Krug, M.; Schwanke, B.; Frey, U.; Kuhl, D. Somatodendritic Expression of an Immediate Early Gene Is Regulated by Synaptic Activity. Proc. Natl. Acad. Sci. U. S. A. 1995, 92 (12), 5734–5738. https://doi.org/10.1073/pnas.92.12.5734.

(3) Epstein, I.; Finkbeiner, S. The Arc of Cognition: Signaling Cascades Regulating Arc and Implications for Cognitive Function and Disease. Semin. Cell Dev. Biol. 2018, 77, 63–72. https://doi.org/10.1016/j.semcdb.2017.09.023.

(4) Zhang, H.; Bramham, C. R. Arc/Arg3.1 Function in Long-term Synaptic Plasticity: Emerging Mechanisms and Unresolved Issues. Eur. J. Neurosci. 2020, ejn.14958. https://doi.org/10.1111/ejn.14958.

(5) Na, Y.; Park, S.; Lee, C.; Kim, D. K.; Park, J. M.; Sockanathan, S.; Huganir, R. L.; Worley, P. F. Real-Time Imaging Reveals Properties of Glutamate-Induced Arc/Arg 3.1 Translation in Neuronal Dendrites. Neuron 2016, 91 (3), 561–573. https://doi.org/10.1016/j.neuron.2016.06.017.

(6) Korb, E.; Wilkinson, C. L.; Delgado, R. N.; Lovero, K. L.; Finkbeiner, S. Arc in the Nucleus Regulates PML-Dependent GluA1 Transcription and Homeostatic Plasticity. Nat. Neurosci. 2013, 16 (7), 874–883. https://doi.org/10.1038/nn.3429.

(7) Chowdhury, S.; Shepherd, J. D.; Okuno, H.; Lyford, G.; Petralia, R. S.; Plath, N.; Kuhl, D.; Huganir, R. L.; Worley, P. F. Arc/Arg3.1 Interacts with the Endocytic Machinery to Regulate AMPA Receptor Trafficking. Neuron 2006, 52 (3), 445–459. https://doi.org/10.1016/j.neuron.2006.08.033.

(8) Wall, M. J.; Corrêa, S. A. L. The Mechanistic Link between Arc/Arg3.1 Expression and AMPA Receptor Endocytosis. Semin. Cell Dev. Biol. 2018, 77, 17–24. https://doi.org/10.1016/j.semcdb.2017.09.005.

(9) Wee, C. L.; Teo, S.; Oey, N. E.; Wright, G. D.; VanDongen, H. M. A.; VanDongen, A. M. J. Nuclear Arc Interacts with the Histone Acetyltransferase Tip60 to Modify H4K12 Acetylation. eNeuro 2014, 1 (1), ENEURO.0019-14.2014. https://doi.org/10.1523/ENEURO.0019-14.2014.

(10) Leung, H. W.; Foo, G. W. Q.; VanDongen, A. M. J. Arc Regulates Transcription of Genes for Plasticity, Excitability and Alzheimer’s Disease. bioRxiv 2019, 833988. https://doi.org/10.1101/833988.

(11) Myrum, C.; Baumann, A.; Bustad, H. J.; Flydal, M. I.; Mariaule, V.; Alvira, S.; Cuéllar, J.; Haavik, J.; Soulé, J.; Valpuesta, J. M.; et al. Arc Is a Flexible Modular Protein Capable of Reversible Self-Oligomerization. Biochem. J. 2015, 468 (1), 145–158. https://doi.org/10.1042/BJ20141446.

(12) Byers, C. E.; Barylko, B.; Ross, J. A.; Southworth, D. R.; James, N. G.; Taylor, C. A.; Wang, L.; Collins, K. A.; Estrada, A.; Waung, M.; et al. Enhancement of Dynamin Polymerization and GTPase Activity by Arc/Arg3.1. Biochim. Biophys. Acta - Gen. Subj. 2015, 1850 (6), 1310–1318. https://doi.org/10.1016/j.bbagen.2015.03.002.

(13) Pastuzyn, E. D.; Day, C. E.; Kearns, R. B.; Kyrke-Smith, M.; Taibi, A. V.; McCormick, J.; Yoder, N.; Belnap, D. M.; Erlendsson, S.; Morado, D. R.; et al. The Neuronal Gene Arc Encodes a Repurposed Retrotransposon Gag Protein That Mediates Intercellular RNA Transfer. Cell 2018, 172 (1–2), 275–288.e18. https://doi.org/10.1016/j.cell.2017.12.024.

(14) Ashley, J.; Cordy, B.; Lucia, D.; Fradkin, L. G.; Budnik, V.; Thomson, T. Retrovirus-like Gag Protein Arc1 Binds RNA and Traffics across Synaptic Boutons. Cell 2018, 172 (1–2), 262–274.e11. https://doi.org/10.1016/j.cell.2017.12.022.

(15) Bloomer, W. A. C.; VanDongen, H. M. A.; VanDongen, A. M. J. Activity-Regulated Cytoskeleton-Associated Protein Arc/Arg3.1 Binds to Spectrin and Associates with Nuclear Promyelocytic Leukemia (PML) Bodies. Brain Res. 2007, 1153 (1), 20–33. https://doi.org/10.1016/j.brainres.2007.03.079.

(16) Digman, M. A.; Brown, C. M.; Sengupta, P.; Wiseman, P. W.; Horwitz, A. R.; Gratton, E. Measuring Fast Dynamics in Solutions and Cells with a Laser Scanning Microscope. Biophys. J. 2005, 89, 1317–1327.

(17) Anton, H.; Taha, N.; Boutant, E.; Richert, L.; Khatter, H.; Klaholz, B.; Rondé, P.; Réal, E.; De Rocquigny, H.; Mély, Y. Investigating the Cellular Distribution and Interactions of HIV-1 Nucleocapsid Protein by Quantitative Fluorescence Microscopy. PLoS One 2015, 10 (2), e0116921. https://doi.org/10.1371/journal.pone.0116921.

(18) Digman, M. A.; Dalal, R.; Horwitz, A. F.; Gratton, E. Mapping the Number of Molecules and Brightness in the Laser Scanning Microscope. Biophys. J. 2008, 94 (6), 2320–2332. https://doi.org/10.1529/biophysj.107.114645.

(19) Phillips, G. N. Structure and Dynamics of Green Fluorescent Protein. Curr. Opin. Struct. Biol. 1997, 7 (6), 821–827. https://doi.org/10.1016/S0959-440X(97)80153-4.

(20) Goo, B. M. S. S.; Sanstrum, B. J.; Holden, D. Z. Y.; Yu, Y.; James, N. G. Arc/Arg3.1 Has an Activity-Regulated Interaction with PICK1 That Results in Altered Spatial Dynamics. Sci. Rep. 2018, 8 (1). https://doi.org/10.1038/s41598-018-32821-4.

(21) Wallrabe, H.; Periasamy, A. Imaging Protein Molecules Using FRET and FLIM Microscopy. Curr. Opin. Biotechnol. 2005, 16 (1 SPEC. ISS.), 19–27. https://doi.org/10.1016/j.copbio.2004.12.002.

(22) Malacrida, L.; Ranjit, S.; Jameson, D. M.; Gratton, E. The Phasor Plot: A Universal Circle to Advance Fluorescence Lifetime Analysis and Interpretation. Annu. Rev. Biophys. 2021, 50, 575–593. https://doi.org/10.1146/annurev-biophys-062920-063631.

(23) Brand, P.; Lenser, T.; Hemmerich, P. Assembly Dynamics of PML Nuclear Bodies in Living Cells. PMC Biophys. 2010, 3 (1), 3. https://doi.org/10.1186/1757-5036-3-3.

(24) Malacrida, L.; Hedde, P. N.; Ranjit, S.; Cardarelli, F.; Gratton, E. Visualization of Barriers and Obstacles to Molecular Diffusion in Live Cells by Spatial Pair-Cross-Correlation in Two Dimensions. Biomed. Opt. Express 2018, 9, 303–321.

(25) Carmichael, R. E.; Henley, J. M. Transcriptional and Post-Translational Regulation of Arc in Synaptic Plasticity. Semin. Cell Dev. Biol. 2018, 77, 3–9. https://doi.org/10.1016/j.semcdb.2017.09.007.

