## Supplement for "Differential mobility and self-association of Arc/Arg3.1 in the cytoplasm and nucleus of living cells"

[1]Beckman Laser Institute and Medical Clinic, [2]Department of Pharmaceutical Sciences, and [3]Laboratory for Fluorescence Dynamics, University of California, Irvine CA 92697. [4]Department of Cell and Molecular Biology, John A. Burns School of Medicine, 651 Ilalo St., BSB 222, University of Hawaii, Honolulu, HI 96813. [5]Department of Pharmacology, University of Texas Southwestern Medical Center, 6001 Forest Park, Dallas, Texas 75390.

### Supplementary Information

#### Supplementary Figure Legends

**Supplementary Figure S1. EGFP-Arc expression levels for N&B analysis.** For each cell included in the N&B analyis shown in Figure 2D, the average EGFP-Arc concentration was calculated based on a reference measurement with an EGFP solution of known concentration. For all cells quantified with N&B, the average expression level of EGFP-Arc was less than 300 nM, a regime in which EGFP does not aggregate.

**Supplementary Figure S2. Fluorescence lifetime imaging of EGFP co-expressed with mCherry.** Cumulative phasor histogram (left) of EGFP fluorescence lifetimes. Before lifetime imaging, single channel images were taken to confirm co-expression of EGFP (right top row) and mCherry (right middle row). In the phasor histogram, pixels corresponding to a single exponential decay of 3.4 ns were selected (cyan circle/cursor) and painted in the corresponding intensity images (right bottom row). No energy transfer between EGFP and mCherry was detected as no shorter, multicomponent lifetimes (selected by magenta circle/cursor) were found. Scale bars, 20 µm.

**Supplementary Figure S3. Fluorescence lifetime imaging of PML-EGFP co-expressed with Arc-mCherry.** (A) Cumulative phasor histogram of PML-EGFP fluorescence lifetimes when co-expressed with Arc-mCherry (before lifetime imaging, single channel images were taken to confirm co-expression of both proteins). In the phasor histogram, pixels corresponding to a single exponential decay of 3.4 ns were selected (cyan circle/cursor) and painted in the corresponding intensity images (B). No energy transfer between PML-EGFP and Arc-mCherry was detected as no shorter, multicomponent lifetimes (selected by magenta circle/cursor) were found. (C) Cumulative phasor histogram of EGFP fluorescence lifetimes in cells expressing PML-EGFP only. As expected for this negative control, no shorter lifetimes were detected (D). Scale bar, 10 µm.

**Supplementary Figure S1**

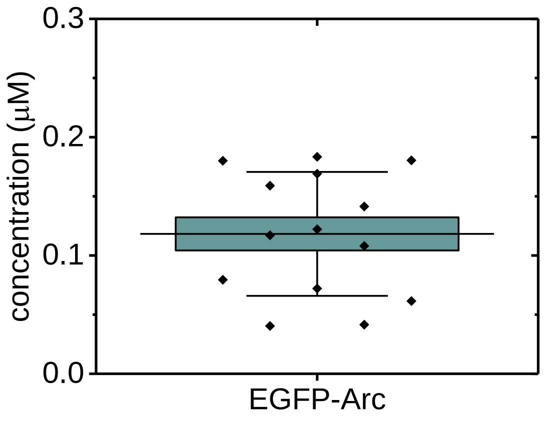

**Supplementary Figure S2**

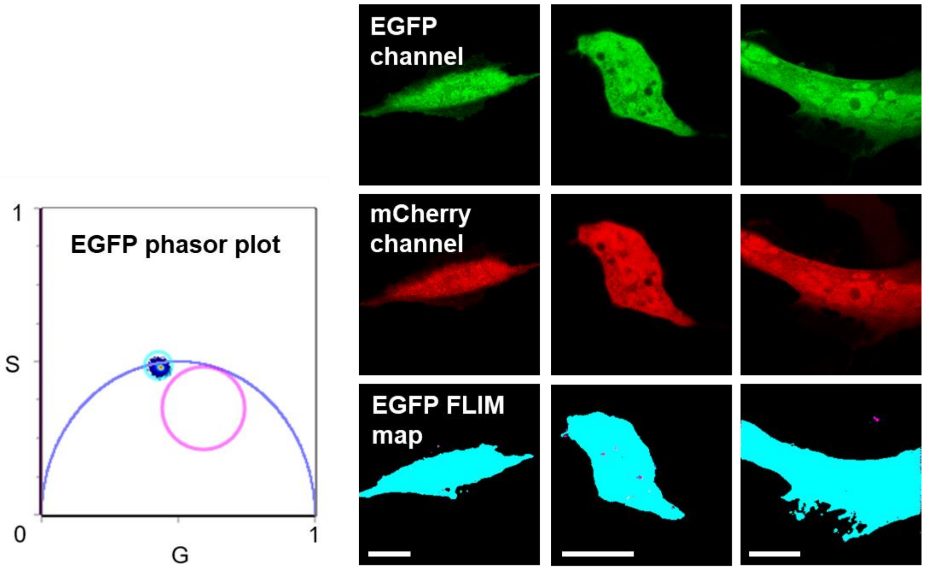

**Supplementary Figure S3**

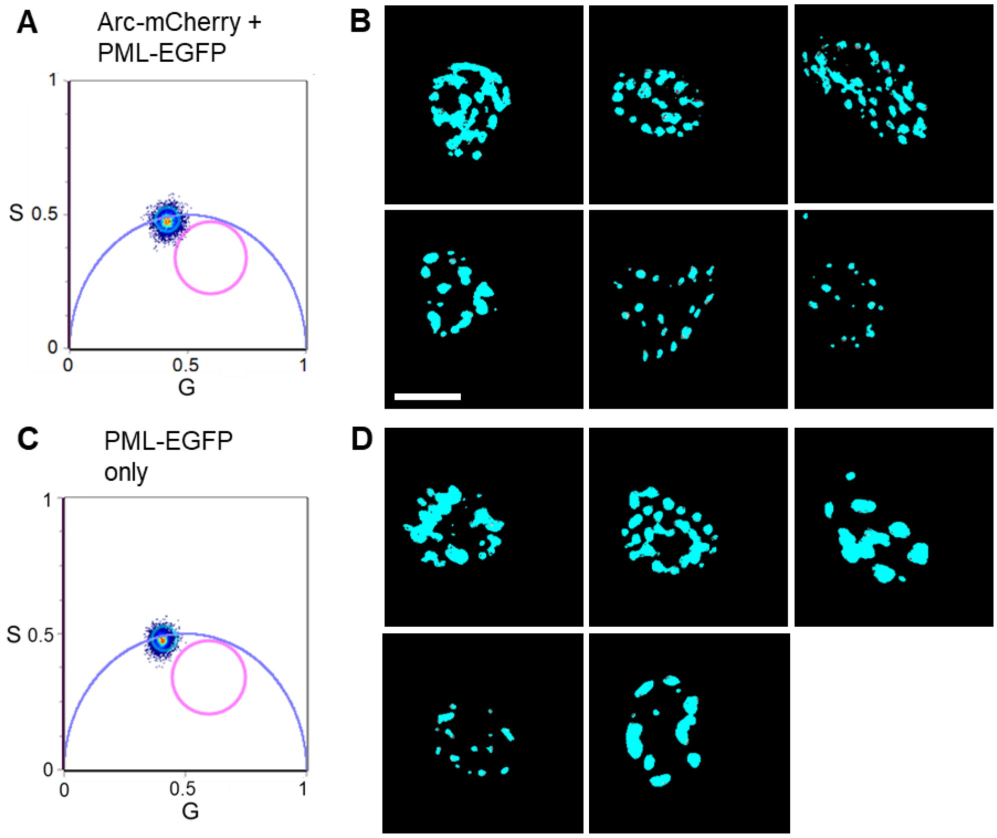
